# Metabolic overflow prevents ecological collapse in microbial populations

**DOI:** 10.1101/2025.06.26.661698

**Authors:** Clemente F. Arias, Francisco J. Acosta, Federica Bertocchini, Cristina Fernandez-Arias

## Abstract

Metabolic overflow is defined by the excretion of partially oxidized byproducts, such as acetate, ethanol, and lactate, which could otherwise be used to produce energy. This behavior, widespread across diverse biological systems, is paradoxical because it suggests a seemingly wasteful use of resources that could instead be employed to maximize energy production. In this work, we investigate the ecological implications of metabolic overflow, using bacterial acetate metabolism as a model. We formulate a second-order generalization of the logistic growth equation, incorporating resource availability and acceleration dynamics to provide an explicit mechanistic link between metabolic overflow and ecological collapse. We demonstrate that acetate-mediated growth deceleration reduces the risk of overshoot and collapse, especially under conditions of resource oversupply. By inhibiting excessive proliferation, acetate overflow enhances the survival and stability of bacterial communities in fluctuating environments. In this context, acetate acts as a quorum-sensing signal, coordinating population-wide responses and enabling the active regulation of growth rates before resource exhaustion occurs. Our findings suggest that overflow metabolism is an adaptive strategy evolved to mitigate the risk of ecological collapse and stabilize microbial populations. These insights highlight the dual role of metabolic byproducts as both inhibitors of growth and regulators of population dynamics, with significant implications for microbial ecology and biotechnological applications.

## Introduction

Metabolic overflow is defined as the excretion of metabolic byproducts that cells could otherwise use to produce energy [1, 2]. This phenomenon is commonly observed in microbial systems like *Escherichia coli* and *Saccharomyces cerevisiae* when oxygen [3] and nutrients, particularly glucose, are abundant [4–7]. Under such conditions, fast-growing cells excrete partially oxidized byproducts (acetate and ethanol, respectively) that could be further metabolized for energy production. A similar behavior is observed in mammalian cells, which excrete lactate in response to heightened metabolic activity, as occurs in immune cells during activation or in rapidly proliferating cells, such as tumor cells [8–10].

Diverting metabolic substrates toward excretion instead of fully utilizing them for energy production is a puzzling behavior. Furthermore, the byproducts of overflow metabolism can negatively impact the cellular function when they accumulate in high concentrations. For instance, acetate can be toxic and disrupts cellular processes by inhibiting key metabolic enzymes, thus retarding bacterial growth [11–13]. Similarly, in yeasts, excess ethanol impairs cell function and inhibits proliferation [13–18]. In mammalian cells, high lactate concentrations contribute to extracellular acidification, which can interfere with cellular processes. Therefore, the excretion of metabolic byproducts is not only seemingly wasteful but also potentially harmful. Despite this, metabolic overflow is observed across a wide range of cellular systems.

The widespread occurrence of overflow metabolism across diverse organisms underscores its fundamental relevance. Bacteria are an ideal model for exploring the causes and consequences of this phenomenon due to their well-characterized biochemical networks and relevance in both industrial and clinical contexts. In biotechnological applications, for instance, acetate is the primary by-product of *E. coli* processes, even under aerobic conditions [11], and it is widely recognized as one of the major constraints limiting biomass and product yields [19, 20].

Considering the detrimental effects of acetate on population growth, it is natural to question why bacteria have not evolved mechanisms to mitigate the consequences of overflow metabolism or prevent its occurrence altogether. The remarkable prevalence of acetate overflow among bacterial species [21–23] suggests that it is deeply ingrained in bacterial physiology. This perspective is further supported by the limited success of engineering efforts aimed at overcoming acetate overflow in biotechnology, despite decades of intensive research [19].

The accumulation of extracellular acetate interferes with bacterial growth at concentrations as low as 0.5 g/L [24]. As a weak acid, acetate contributes to medium acidification and cellular toxicity. However, its effects on bacterial growth extend beyond toxicity, and point to more subtle and complex regulatory roles for acetate overflow [12]. It has recently been shown that acetate is a global regulator of glucose metabolism in *E. coli*, inhibiting the expression of multiple genes involved in glucose uptake, glycolysis, and the TCA cycle [12, 25]. Consequently, acetate reduces both the production of acetyl-CoA from glucose and its use to generate energy or synthesize anabolic precursors for growth [12].

The control of acetyl-CoA production and consumption by acetate provides a mechanistic explanation for the inhibitory effect of overflow metabolism on growth rates [12]. Acetate freely permeates the cell membrane, and under overflow conditions, high extracellular acetate concentrations cause a substantial inflow into the cytoplasm [26]. This intracellular accumulation of acetate may dampen metabolic activity within individual cells, which translates into a reduction in population growth.

Regardless of their precise mechanistic connection, acetate overflow and growth rates are linked through a negative feedback loop. Empirical evidence shows that growth rates dictate the dynamics of acetate excretion according to a well-defined threshold-linear pattern [5, 20, 27]. In turn, acetate accumulation inhibits growth in a concentration-dependent manner [13] (Fig. 1).

**Figure 1.**
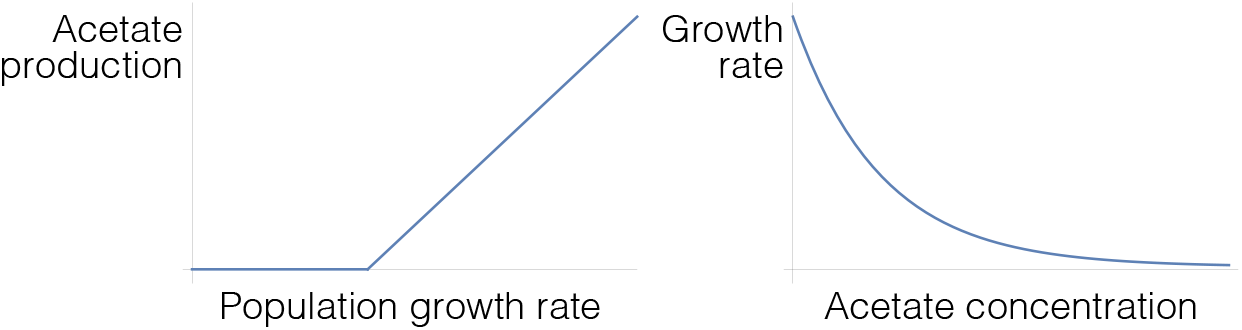
Population-level feedback loop between growth rates and acetate overflow. A) Threshold-linear relationship between acetate production and growth rate: acetate excretion remains negligible below a certain threshold but increases linearly once this threshold is exceeded [5, 20, 27]. B) Acetate, in turn, reduces growth rates in a dose-dependent manner [13].

In this work, we propose that this feedback loop decelerates growth, particularly in rapidly proliferating populations. We further suggest that acetate-mediated growth deceleration mitigates the risk of ecological collapse in bacterial populations. Collapse often occurs when a population exceeds the capacity of the environment to sustain growth—a phenomenon known as overshoot [28, 29]. Excessive growth can deplete available resources, triggering a rapid contraction that may lead to extinction. We hypothesize that bacterial populations use extracellular acetate concentrations as a signal to monitor growth. High acetate levels indicate rapid proliferation, a situation likely to result in overshoot and collapse. By using acetate to decelerate growth, bacterial populations could avoid this outcome. From this perspective, acetate serves as a regulatory signal that modulates proliferation rather than acting as a toxin, providing an effective mechanism to prevent extinction and enhance the chances of survival.

To assess the validity and implications of this hypothesis, we adopt a modeling approach. Mathematical models have been widely used to investigate various aspects of metabolic overflow [26, 30–35]. In this work, we focus on the effects of overflow metabolism on population dynamics. We show that the classical logistic growth equation can be generalized to model overshoot and ecological collapse. Using this extended model, we illustrate how a transient increase in resources can lead to excessive growth, overshoot, and ultimately extinction. Then, we demonstrate that metabolic overflow effectively mitigates this risk by decelerating growth, helping to stabilize the population and prevent ecological collapse. Finally, we show that the strategy of growth deceleration provides competitive advantage over populations lacking this mechanism and may be crucial for the long-term survival of microbial communities.

This work suggests a novel adaptive role for metabolic overflow in population dynamics. By showing that growth deceleration can prevent ecological collapse, we propose a mechanism that helps to stabilize microbial populations after the colonization of new environments or under unexpected resource fluctuations. These findings provide fresh insights into microbial ecology and open new directions for improving biotechnological processes where managing growth and overflow metabolism is essential.

## Results

### Second-order models reproduce ecological overshoot and collapse

Biological systems cannot grow indefinitely. The existence of an upper limit to size was already explicit in the earliest mathematical formalizations of biological growth [36, 37]. In classical population dynamics models, this limit takes the form of a predefined parameter. For instance, logistic growth is modeled by the following equation:

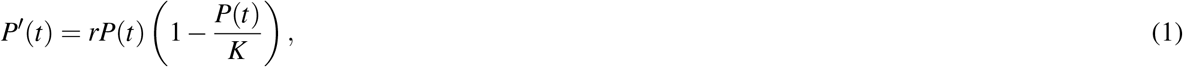

where *P*(*t*) is the population size at time *t, r* represents the population’s intrinsic growth rate, and *K* is the asymptotic size of the system, typically referred to as the carrying capacity.

External constraints imposed by the environment are inherent to biological growth and exist independently of any specific modeling framework. However, the logistic model assumes that these constraints gradually limit proliferation as population size increases. In reality, environmental constraints often deviate from this pattern, and living systems can transiently exceed their carrying capacity—a phenomenon known as overshoot [28, 45]. Such transient dynamics, which are widely observed in natural systems ranging from ecological populations [38, 39] to tumors [40–42] and stem cells [43], cannot be adequately captured by the logistic framework [28, 44–47].

In this work, we propose that overflow metabolism plays a crucial role in preventing overshoot and collapse in microbial populations. Testing this hypothesis requires an alternative modeling framework capable of capturing overshoot dynamics. One such framework was introduced by Hutchinson [48, 49], who modified the logistic equation to account for delayed density-dependent effects on population growth:

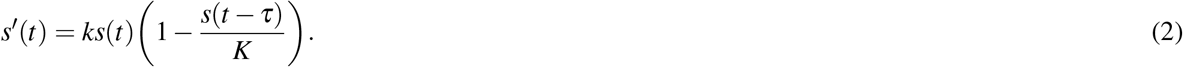

This model assumes that the constraints imposed by the carrying capacity do not immediately affect the population’s size but instead act with a certain delay *τ* [50].

Equation 2 is a phenomenological model originally intended to reproduce the cyclic oscillations observed in certain ecological populations [48, 49]. Additionally, it predicts population collapse as a possible outcome of overshoot under specific combinations of parameter values (Fig. 2.A).

**Figure 2.**
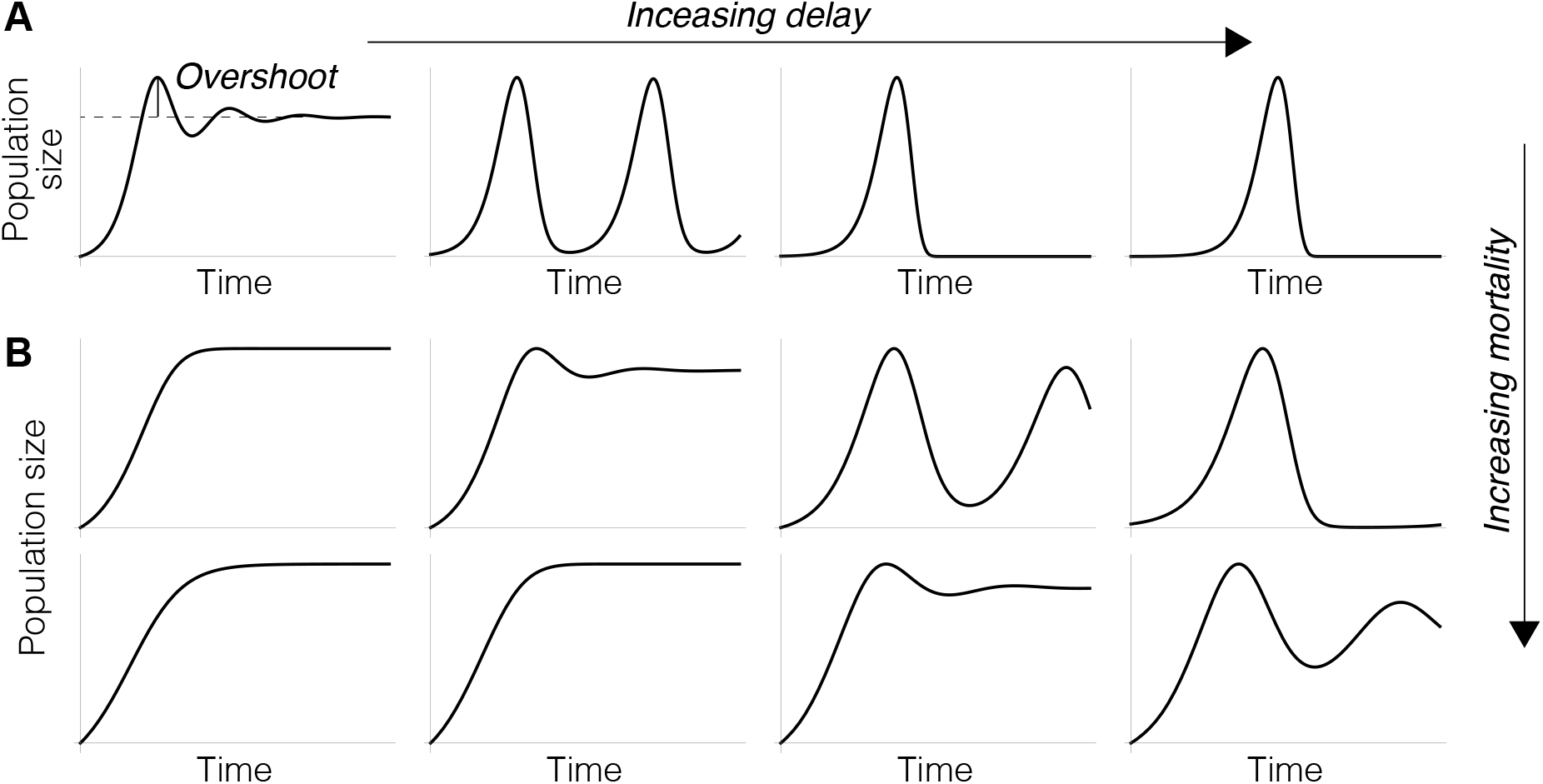
Dynamics of models 2 and 3. A) Model 2 captures overshoot and oscillatory dynamics, ultimately leading to collapse as the time delay (*τ*) increases. B) Model 3 predicts that increasing mortality (parameter *µ*) suppresses oscillations and prevents collapse. Parameter values: *r* = 0.5, *K* = 3, *P*_0_ = 0.1 for *τ* = 2, 4, 8, 12. B) *µ* = 0.3 and 0.4.

According to this equation, the intensity and consequences of overshoot only depend on delay, with shorter delays resulting in lower overshoot and longer delays leading to collapse (Fig. 2.A). However, in real populations, such extreme outcomes are unlikely to arise from variations in the timing of density-dependent effects alone. This is evident in populations that exhibit overshoot under specific conditions but not in others. A classic example is the reindeer population introduced to the St. Matthew Island in 1944. The absence of predators and the abundance of resources allowed the population to grow exponentially, eventually leading to overshoot and collapse as resources were depleted [51]. In contrast, in environments where natural predators are present, reindeer population growth is constrained, effectively preventing overshoot.

Equation 2 can be modified to reproduce these mortality-mediated dynamics:

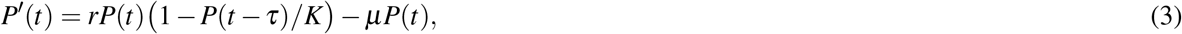

where *µ* represents the mortality rate due to factors unrelated to density-dependent effects. Variations in this parameter can determine whether the population survives or becomes extinct after overshoot (Fig. 2.B).

This model emphasizes the role of mortality in preventing overshoot and collapse. However, increased mortality is not the primary effect of overflow metabolism. Although acetate can cause cell death in bacterial populations, particularly at high concentrations, its main effect is growth inhibition [13]. As discussed earlier, acetate decelerates proliferation by reducing glucose uptake and limiting anabolic processes in bacterial cells [11–15]. Consequently, acetate does not directly alter the population size but instead reduces its growth rate. We propose that this acetate-induced deceleration is better captured through a second-order differential equation framework. Specifically, we suggest the following second-order generalization of the logistic growth model:

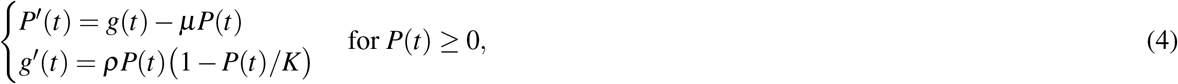

where *K* is the carrying capacity and *ρ* denotes the population’s intrinsic acceleration rate, i.e. the maximum rate at which the population’s growth rate increases under optimal, ideal conditions.

This model distinguishes between first-order effects, such as mortality, which directly change the population size, and second-order effects, which modify the population’s growth rate. Specifically, the density-dependent effects resulting from the limited carrying capacity of the environment are reflected in changes to the acceleration of population size.

To understand how equations 4 extend the classical logistic growth model, let us rewrite them as

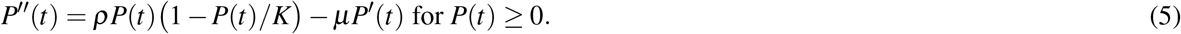

This expression reveals that model 4 approaches the logistic equation with intrinsic growth rate *ρ/µ* as parameters *ρ* and *µ* tend to infinity (Fig. 3.A).

**Figure 3.**
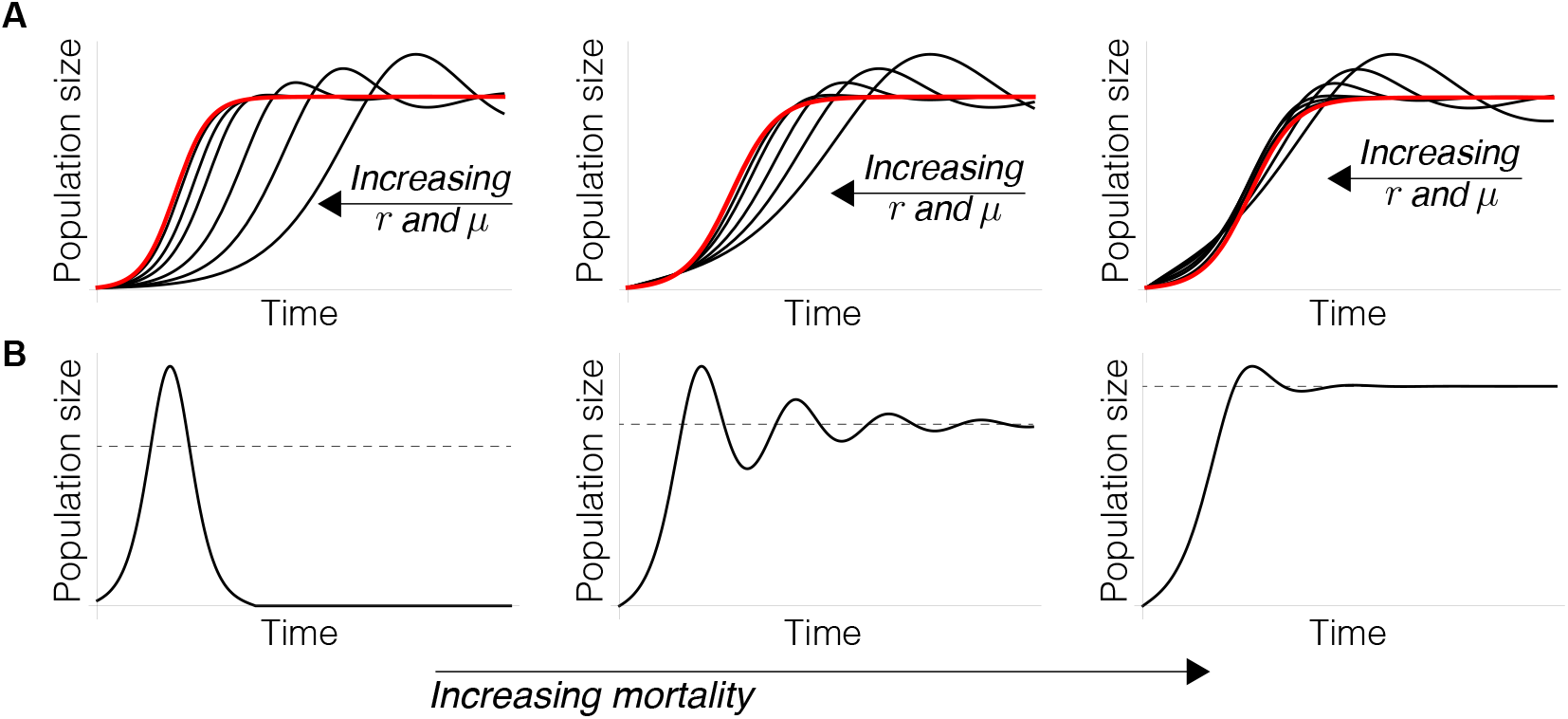
Dynamics of model 4. A) The model can be interpreted as a generalization of classical logistic growth. As the parameters *ρ* and *µ* increase, the model behavior converges to that of the logistic equation (orange lines). B) The model reproduces collapse under certain conditions (left), but this effect is mitigated as mortality rates increase (center and right). Parameter values: A) *r* = 0.3, 0.6, 1.2, 3, 6, 24, *K* = 10, *P*_0_ = 0.1, *g*_0_ = 0 (left), 0.4 (center),1 (right), *µ* = 0.25, 0.5, 1, 2.5, 5, 20. B) *r* = 0.5, *K* = 3, *P*_0_ = 0.1, *g*_0_ = 0.1, and *µ* = 0, 0.2, 0.6.

Second-order differential equations have been previously used to model population dynamics in the literature [50, 52]. Our choice of Model 5 is justified because it naturally extends the logistic growth model and captures behaviors such as overshoot, collapse, and the impact of increased mortality rates on population survival (Fig. 3.B). In the following sections, we will show that incorporating the acceleration of the population’s size provides an effective framework for modeling deceleration, which is the primary effect of overflow metabolism on microbial population dynamics.

### Oversupply of resources may lead to overshoot and extinction

The notion of carrying capacity, originally defined as the upper limit on the size of growing natural systems [28, 29, 53], is crucial for understanding overshoot. This concept is widely used across biological disciplines to describe the interactions between populations and their environment [29]. However, the carrying capacity lacks a rigorous formalization and is often used ambiguously to summarize the effect of environmental constraints on growth [54]. Moreover, it is typically assumed to be a static parameter, implying that these constraints remain constant despite potentially changing conditions [29, 53]. In reality, the maximum population size that a given environment can sustain is dynamic and may fluctuate, for instance, with variations in resource availability or quality [50].

To address these limitations, we propose a more mechanistic definition of carrying capacity by explicitly linking it to resource availability. To this end, we extend model 4 to include the dynamics of a specific resource, *R*:

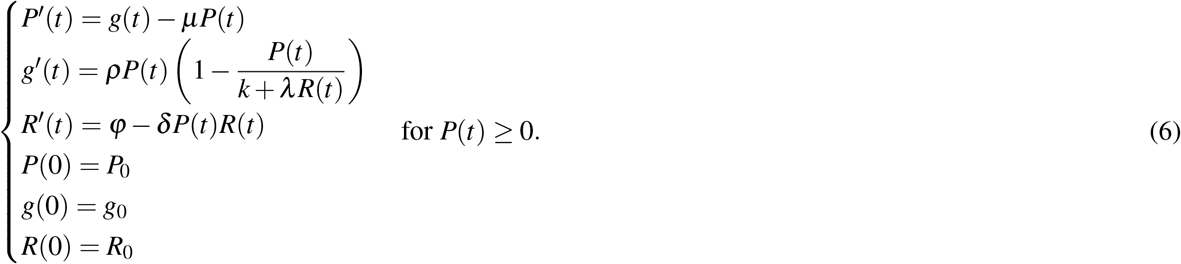

where *R*(*t*) represents the availability of resource *R* at time *t*, and *f* (*t*) *>* 0 and *δ >* 0 represent the inflow of *R* into the system and the per-capita rate of resource consumption by the population, respectively.

In this model, parameter *K*, which denotes the carrying capacity in the logistic equation (Equation 4), is replaced by the expression:

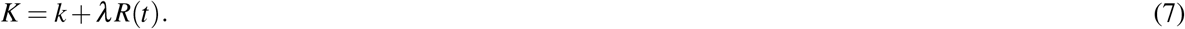

Parameter *k* represents a baseline carrying capacity that is independent of *R*. If *R* is the only resource driving population growth, then *k* = 0. Conversely, if *k >* 0, the population can sustain itself by consuming other resources not explicitly considered in the model. Parameter *λ*, on the other hand, quantifies the contribution of the resource *R* to the overall carrying capacity. A higher value of *λ* implies that *R* has a stronger effect on increasing the carrying capacity of the system, whereas a lower value indicates a weaker impact of *R*.

Equation 6 provides a general framework for understanding how the dynamics of specific resources contribute to the onset of overshoot. To illustrate this point, let us consider a situation where *f* (*t*) = *ϕ*, a positive constant. Under this condition, the system attains a steady state given by:

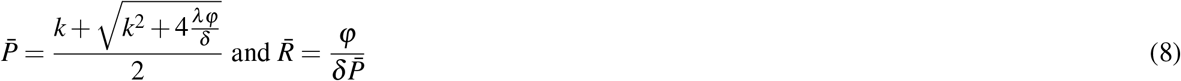

The steady-state population size, 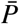, depends on the ratio between the rates of resource supply (*ϕ*) and resource consumption (*δ*). This expression underscores a critical aspect of biological growth that is only implicitly captured in the concept of carrying capacity: populations are open systems that must consume resources to survive. Population growth can only be sustained if this consumption is balanced by the continuous replenishment of resources. This becomes particularly evident when *k* = 0 in model 6, which implies that *R* is the sole limiting factor for population growth. In this case, the population only survives in the long term if there is a sustained input of resources (i.e. if *ϕ >* 0).

In the previous section, we showed that equations 4 approach the classical logistic model for large values of the intrinsic acceleration rate *ρ* and the mortality rate *µ*. In this case, 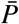 coincides with the upper limit of the population size (Fig. 4.A). However, in other scenarios, the system can exceed this steady-state value (Fig. 4.B). In this work, we define overshoot as the transient condition in which the population temporarily exceeds its long-term carrying capacity.

**Figure 4.**
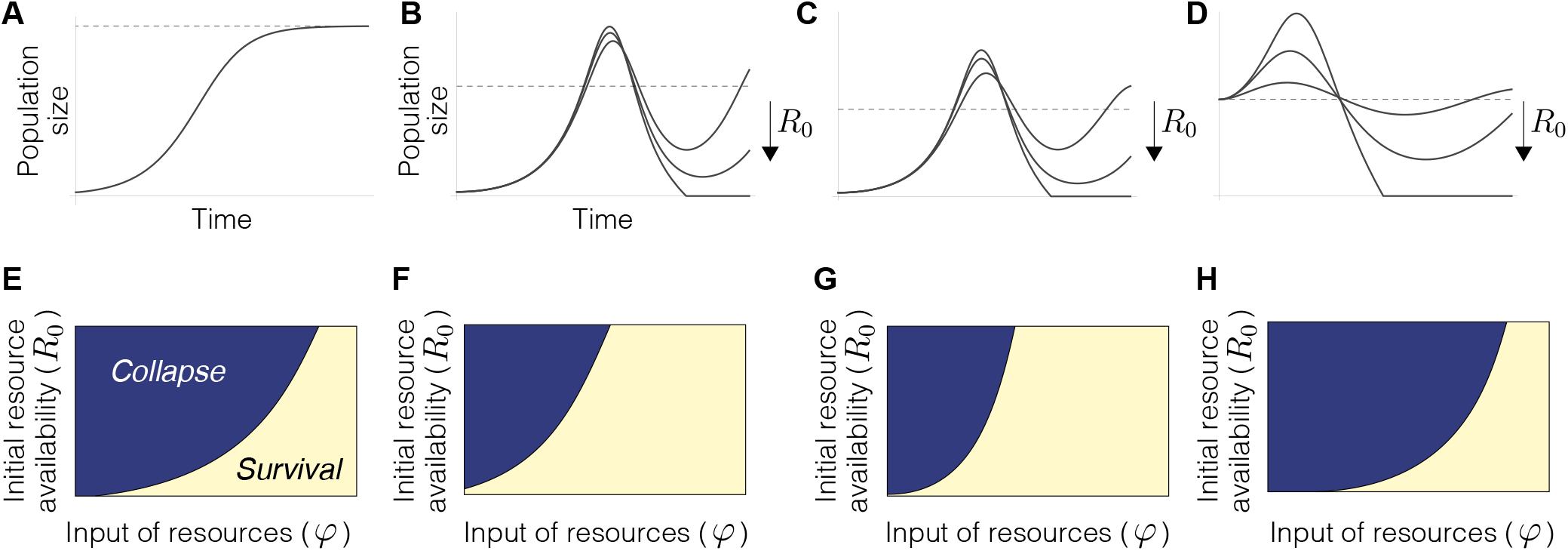
Dynamics of Model 6. A) For large values of the parameters *ρ* and *µ*, the population approaches its asymptotic value without overshoot. B) For lower values of these parameters, the population exhibits overshoot. During the colonization of new environments, greater resource availability (*R*_0_) increases the extent of overshoot and may lead to collapse. C) Collapse can occur even if resource *R* is not the sole determinant of the carrying capacity (*k >* 0). D) Stable populations may also collapse after a sudden pulse of resources. E) In the colonization of new environments, the risk of collapse increases with higher resource availability (*R*_0_) and lower resource input (*ϕ*). F) Lower initial growth rates (*g*_0_) reduce the risk of collapse in these circumstances. G) Similar outcomes are observed for stable populations after a pulse of resources. H) The risk of collapse increases when the contribution of *R* to the carrying capacity is greater (corresponding to lower values of parameter *k*). Parameter values: A) *µ* = 10, *ρ* = 10, *k* = 5, *λ* = 0.4, *ϕ* = 1, *δ* = 0.5, *P*_0_ = 0.1, *g*_0_ = 0, *R*_0_ = 5. B) *µ* = 0.1, *ρ* = 0.1, *k* = 0, *ϕ* = 9, *R*_0_ = 10, 400, 900. C) *k* = 1, *ϕ* = 4. D)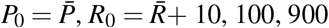. E) 0 ≤ *ϕ* ≤20, 0 ≤*R*_0_ ≤5000, *g*_0_ = 0.5. Rest of parameters as in B. F) *g*_0_ = 0. Rest of parameters as in E. G) 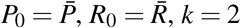. Rest of parameters as in E. H)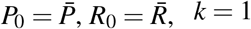. Rest of parameters as in F. Dashed lines in A-D indicate the carrying capacity.

For a given value of resource input (*ϕ*), the magnitude and consequences of overshoot are highly sensitive to initial resource availability (parameter *R*_0_). When resources are initially low, growth is slower, and the population may recover quickly from overshoot. However, when there is an initial surplus of resources, the population can grow rapidly beyond its sustainable limits, potentially leading to collapse or extinction (Fig. 4.B). Importantly, this may occur even if resource *R* is not the only determinant of the population’s carrying capacity (Fig. 4.C). This situation may arise during the colonization of new environments, where nutrients are initially abundant but later decline as the population grows and consumes them. However, such dynamics can also occur in stable populations. A pulse of resources can drive the population away from its steady state, leading to fluctuations that may ultimately result in extinction (Fig. 4.D).

The recovery after overshoot critically depends on the rate at which resources are supplied into the system (parameter *ϕ*). Higher resource input can support recovery, replenishing resources more quickly after the early population boost triggered by initial resource oversupply. In contrast, if the input rate is too low, recovery may be compromised, increasing the risk of collapse for lower initial resource availability (Fig. 4.E). These results suggest that the long-term capacity of an environment to support a population depends on the rate of resource production (*ϕ*). Overshoot, on the other hand, is caused by a transient oversupply of resources.

The consequences of overshoot also depend on the population’s inherent dynamic features. For instance, the population’s initial growth rate (*g*_0_) affects the risk of overshoot and collapse in model 6. Higher values of *g*_0_, reflecting faster initial proliferation, can lead to faster resource depletion and a greater likelihood of the population exceeding sustainable limits, thereby increasing the risk of collapse (Figs. 4.E,F). This parameter can be interpreted as an indicator of adaptation or preconditioning to the environment. A high value of *g*_0_ represents a pre-adapted population, possibly due to prior exposure to similar conditions, which enables rapid growth. Conversely, a lack of preconditioning results in slower initial growth as the population gradually adapts to a new environment. Additionally, *g*_0_ can vary across species with different ecological strategies. For instance, opportunistic species are expected to exhibit higher *g*_0_ values, allowing them to quickly colonize and exploit available resources.

Similar results are observed for stable populations following a pulse of resource *R* (Figs. 4.G). The risk of extinction decreases with the contribution of *R* to the carrying capacity (Figs. 4.G,H). However, these results suggest that a sudden excess of a given resource can destabilize ecological communities, even when the population can grow on additional nutrients. The probability of destabilization and collapse due to a sudden input of secondary metabolites can be mitigated by catabolite repression, a regulatory mechanism that enables microorganisms to prioritize preferred carbon sources, thereby suppressing the expression of genes involved in the metabolism of alternative resources [55].

In this section, we have included the dynamics of resource supply and consumption in equations 4. In this extended model, the carrying capacity is dynamically defined as a function of resource availability. Equations 6 account for the role of resource oversupply in the onset of overshoot and collapse. They also highlight the role of the population’s dynamic parameters in both the colonization of new environments and the stability of already established populations facing a sudden oversupply of resources. In the following sections, we will use this framework to understand the effect of metabolic overflow on population dynamics.

### Metabolic overflow mitigates the risk of ecological collapse

In this section, we will extend equations 6 to include the dynamics of overflow metabolism and its consequences for population dynamics. To do this, we will use the case of acetate overflow and its inhibitory effect on bacterial populations as a reference. Denoting the extracellular concentration of acetate by *A*(*t*), we define the following equations:

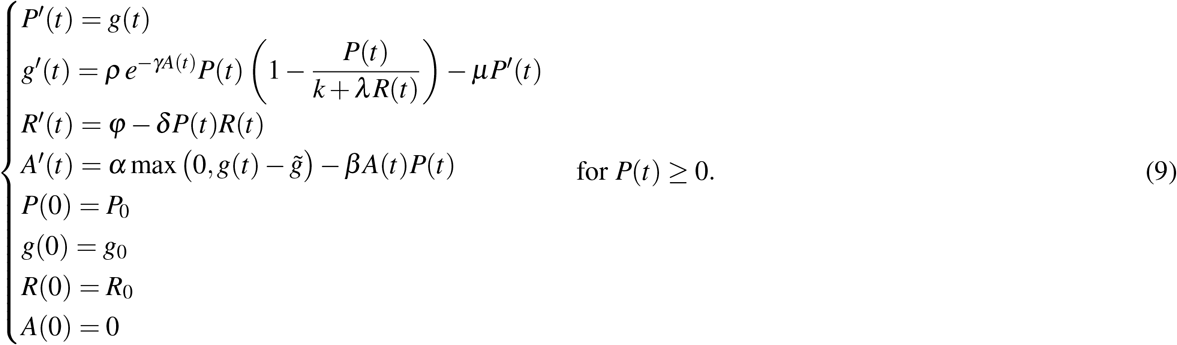

This model assumes that acetate is produced in a threshold-linear manner, depending on the population growth rate (*P*^*′*^(*t*) = *g*(*t*)). Acetate excretion is zero below a threshold rate, 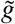, and increases linearly (at a rate *α*) with *g*(*t*) above this threshold. It also assumes that bacterial cells simultaneously excrete and internalize acetate, as empirically observed [12, 19]. The per-capita rate of acetate internalization is denoted by *β*. As for the effects of acetate on population dynamics, it is assumed to reduce the growth rate in a concentration-dependent manner according to the expression *e*^*−γA*(*t*)^.

These assumptions reflect key aspects of the negative feedback loop that links acetate dynamics to growth rates in bacterial populations: acetate excretion increases with population growth [5, 20, 27], while acetate inhibits growth in a concentration-dependent manner [13] (see Fig. 1). The result of this feedback mechanism is a deceleration of bacterial growth, an effect that is naturally modeled by equations 9.

Numerical simulations of these equations show that metabolic overflow significantly reduces the extent of overshoot, an effect that increases with the rate of acetate excretion (parameter *α*) (Fig. 5.A). Growth deceleration caused by acetate overflow may prevent collapse both during the colonization of new environments (Fig. 5.B) and in cases of resource oversupply in stable populations at steady state (Fig. 5.C).

**Figure 5.**
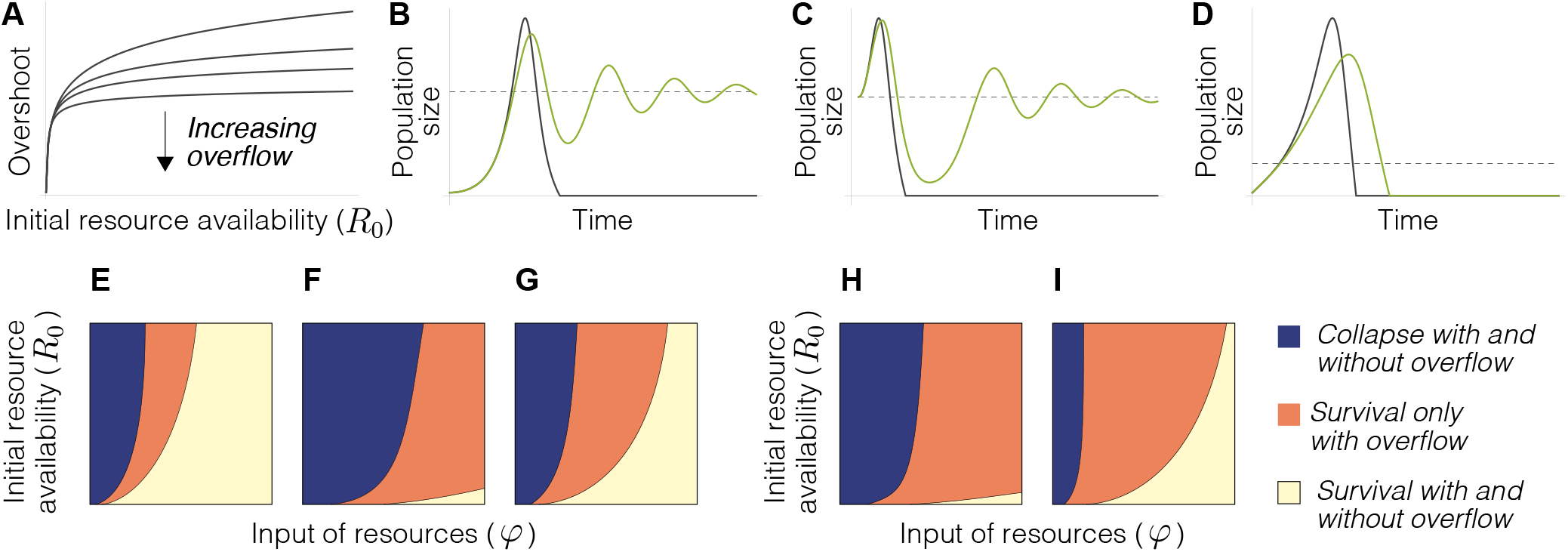
Metabolic overflow mitigates the risk of ecological collapse. A) According to Model 9, overflow reduces the extent of ecological overshoot. B,C) Acetate overflow can prevent collapse both during the colonization of new environments (B) and in stable populations after a pulse of resources (C). D) Metabolic overflow does not always prevent collapse. Lower resource input and greater initial growth rates may lead to population extinction. Green and black lines represent the dynamics of populations with and without metabolic overflow, respectively. Dashed lines indicate the carrying capacity. E-G) Dependence of collapse and survival on resource input (*ϕ*) and initial resource availability (*R*_0_) with and without acetate overflow. Acetate overflow expands the range of environmental conditions in which populations can survive. This effect is more pronounced for greater intrinsic growth rates (F) and higher initial growth rates (G). H,I) Acetate overflow also increases the chances of survival in stable populations after a pulse of resources, an effect that is more pronounced for populations with greater intrinsic growth rates (H). Parameter values: A) *ρ* = 0.4, *γ* = 0.01, *k* = 1, *λ* = 0.4, *µ* = 0.3, *ϕ* = 10, *δ* = 0.5, *α* = 0,100,200,400, 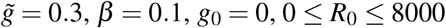 *P*_0_ = 0.1, *R*_0_ = 600, *α* = 0 (black line) and 60 (green line). C) 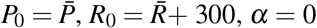 (black line) and 100 (green line). D) *P*_0_ = 0.1, *R*_0_ = 600, *ϕ* = 0.1, *g*_0_ = 0.5, *α* = 0 (black line) and 60 (green line). The parameters not indicated in B-D are as in A. E) *ρ* = 0.2, 0 ≤ *ϕ* ≤ 20, 0 ≤*R*_0_ ≤ 5000, *g*_0_ = 0, *α* = 0 and 100. Rest of parameters as in A. F) *ρ* = 0.4. Rest of parameters as in E. G) *g*_0_ = 0.5. Rest of parameters as in E. H,I) 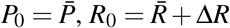, with 0 ≤Δ*R* ≤ 5000, *ρ* = 0.4 (H) and 0.2 (I). Rest of parameters as in E.

The protective effect of overflow metabolism is not universal, and it may not always prevent collapse (Fig. 5.D). However, it significantly increases the range of conditions under which population growth can be sustained. Specifically, populations that exhibit higher initial growth rates (*g*_0_) and greater intrinsic acceleration rates (*ρ*), which are more vulnerable to collapse due to rapid resource depletion and overshoot, tend to benefit more from acetate overflow (Figs. 5.E-I). This protective effect is particularly important when these populations are exposed to environmental conditions that might otherwise lead to unsustainable growth, such as resource oversupply or the colonization of new environments.

In this section, we have extended equations 6 to model metabolic overflow. We have argued that second-order models offer a natural framework for interpreting the deceleration induced by metabolic overflow in population growth. This model suggests that acetate-mediated inhibition of growth rates, particularly in rapidly proliferating populations, may reduce the extent of overshoot and mitigate the risk of subsequent collapse.

### Metabolic overflow is an advantageous strategy in ecological communities

In this section, we will demonstrate that the inhibitory effect of overflow metabolism provides a competitive advantage over populations lacking this mechanism. We will also show that this mechanism can contribute to the long-term survival of bacterial communities. To do this, we will use the following equations:

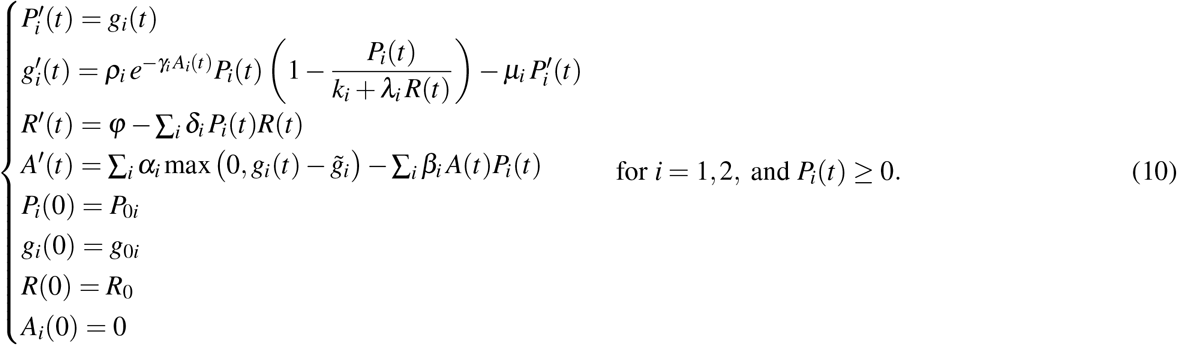

In this model, two populations, *P*_1_ and *P*_2_, compete for resource *R*, consuming it at rates *δ*_1_ and *δ*_2_. According to model 10, both two populations can coexist in the long term. Their respective steady-state sizes are given by

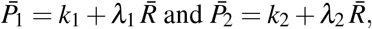

where 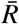 denotes the steady-state concentration of resource *R*, defined as:

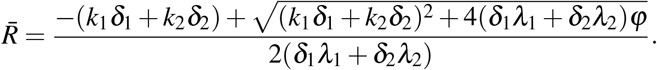

The coexistence of both species does not require metabolic overflow (Fig. 6.A). However, the dynamics of resource *R* can lead to overshoot and collapse (see Figs. 6.B,C and Supplementary Material 1, hybrid systems). According to model 10, overflow metabolism can reduce the extent of overshoot and prevent collapse in microbial communities, thereby facilitating the coexistence of competing populations (Fig. 6.D).

**Figure 6.**
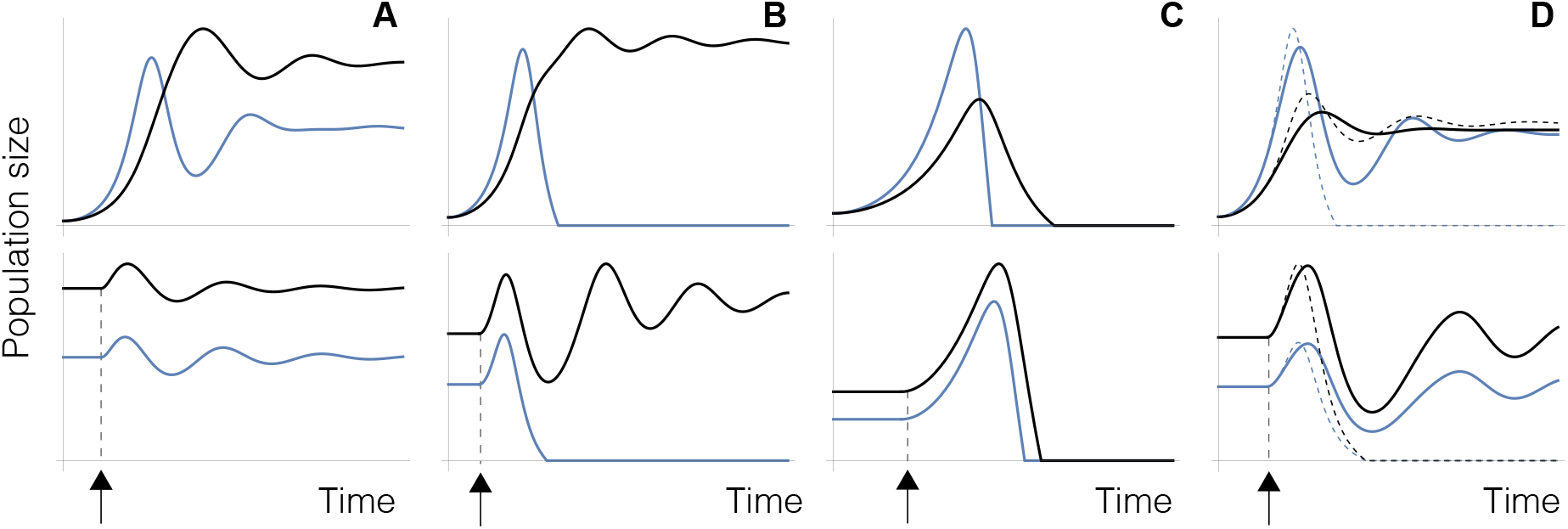
Dynamics of model 10. The figures show the dynamics of two populations during the colonization of new environments (upper) and after a pulse of resources in stable communities (lower). A) Both populations can coexist in the absence of acetate overflow (*α*_*i*_ = 0, *β*_*i*_ = 0, *γ*_*i*_ = 0). B, C) Excess initial resources (*R*_0_) or low resource input (*ϕ*) may lead to the collapse of one (B) or both populations (C). D) Acetate overflow can reduce overshoot and prevent collapse. Solid and dashed lines represent the dynamics of both populations with and without acetate overflow, respectively. Parameter values: A) *R*_0_ = 80, *ϕ* = 20, *ρ*_1_ = 0.15, *µ*_1_ = 0.2, *λ*_1_ = 0.3, *k*_1_ = 0, *δ*_1_ = 0.5, *P*_1_ = 0.1, *r*_2_ = 0.1, *µ*_2_ = 0.2, *λ*_2_ = 0.5, *k*_2_ = 0, *δ*_2_ = 0.5, *P*_2_ = 0.1. (up). 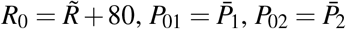(down). B) *R*_0_ = 200, *ϕ* = 5 (up).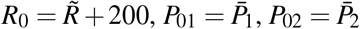(down). C) *R*_0_ = 200, *ϕ* = 0.5 (up). 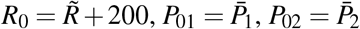 (down). The parameters not indicated in B and C are as in A. D) *R*_0_ = 500. *ϕ* = 10, *r*_1_ = 0.15, *µ*_1_ = 0.2, *λ*_1_ = 0.3, *k*_1_ = 1, *r*_2_ = 0.1, *µ*_2_ = 0.2, *λ*_2_ = 0.5, *k*_2_ = 1, *α*_1_ = *α*_2_ = 200, 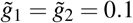(up). 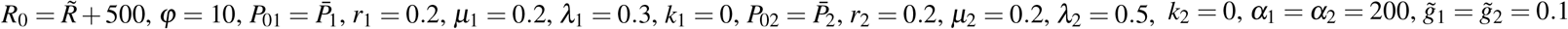 (down).

To further analyze how overflow metabolism affects the risk of overshoot and collapse in the two-population system described by equations 10, we define overflow and non-overflow strategies as follows: A population *P*_*i*_ is said to adopt an overflow strategy if it contributes to the dynamics of extracellular acetate (i.e. *α*_*i*_ *>* 0 and *β*_*i*_ *>* 0) and experiences the inhibitory effect of acetate (*γ*_*i*_ *>* 0). Conversely, the non-overflow strategy is characterized by the absence of acetate secretion and internalization (i.e., *α*_*i*_ = 0 and *β*_*i*_ = 0), as well as the absence of acetate’s effects on population dynamics (*γ*_*i*_ = 0).

Numerical simulations of model 10 show that overflow strategies reduce the risk of population extinction. Collapse during the colonization of new environments is more frequent when both populations adopt non-overflow strategies (Fig. 7.A). In direct competition, overflow strategies significantly improve survival at the expense of non-overflow strategies (scenario S2, Fig. 7.A). Moreover, in communities where both populations employ overflow metabolism (scenario S3, Fig. 7.B), the chances of survival for both populations are greatly enhanced relative to when both populations adopt non-overflow strategies (scenario S1, Fig. 7.B). Similar results are observed in stable communities, where a sudden input of resources can lead to extinction (Figs. 7.C,D). The likelihood of survival increases for populations that adopt the overflow strategy (scenarios S2 and S3, Fig. 7.C). This strategy enhances the simultaneous survival and reduces the simultaneous extinction of both populations (scenarios S3 vs. S1, Fig. 7.D). As shown in Fig. 3.B, higher mortality rates reduce the magnitude of overshoot and, consequently, the risk of ecological collapse. Consistent with this, model 10 predicts that the probability of extinction decreases as mortality rates increase, both during the colonization of new environments (Figs. 7.E,F) and in stable communities (Figs. 7.G,H). Under these conditions, overshoot is less likely to occur (Fig. 3.B), diminishing the differences between overflow and non-overflow strategies. However, overflow strategies remain more advantageous in terms of enhancing both population and community survival.

**Figure 7.**
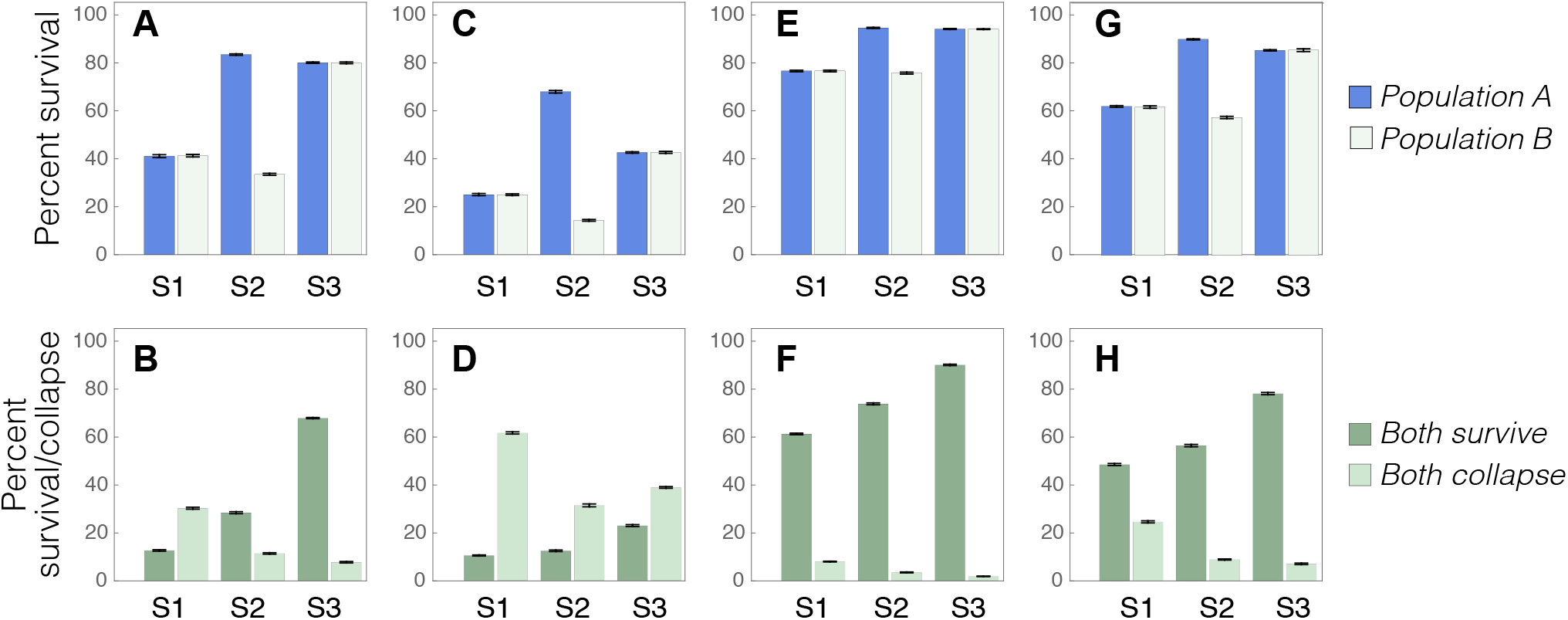
Metabolic overflow confers a competitive advantage. The figures show the probability of survival and collapse of two competing populations under three scenarios: neither population uses overflow strategies (S1), only Population 1 uses this strategy (S2), and both populations use it (S3). A) Probability of survival of both populations during the colonization of a new environment. B) Probability of survival or extinction of both populations under the same conditions. C,D) Same as A,B, but after a pulse of resources in a community at steady state. E-H) Same as A-D, but with higher mortality rates. Error bars represent standard deviations. Details of the simulations and parameter values can be found in Supplementary Material A.

From a game-theory perspective, overflow strategies are evolutionarily stable according to model 10, meaning they cannot be outcompeted by alternative strategies. Cheater populations, for instance, might avoid adopting overflow metabolism to bypass growth deceleration, seemingly gaining a competitive advantage. However, such populations face a greater risk of collapse when excess resources are available. Moreover, in direct competition, populations employing overflow strategies consistently outperform those that do not (scenario S2 in Fig. 7). Consequently, overflow metabolism supports population survival while enhancing the stability and resilience of microbial communities, even in the presence of competitive dynamics.

## Discussion

In this work, we use mathematical modeling to explore the consequences of metabolic overflow on the dynamics of microbial populations. To that end, we introduce a second-order version of the classical logistic growth model. This framework incorporates acceleration, providing a natural way to model acetate-induced deceleration in population growth. The explicit inclusion of resources in our model provides a dynamic definition of the carrying capacity. Furthermore, it highlights the critical role of resource oversupply in triggering overshoot and eventual collapse in ecological populations. In this context, our findings suggest that metabolic overflow significantly enhances the chances of survival for microbial communities facing unexpected fluctuations in resource availability.

Our models focus on the well-established negative feedback loop linking acetate excretion and growth rates, without considering other roles of acetate in bacterial physiology. For example, bacterial cells can use acetate as a carbon source, either simultaneously with glucose or after glucose depletion [1, 6, 19, 26]. The inhibition of growth by acetate [13] suggests that its inhibitory effects are more significant than its potential metabolic contribution in our context, particularly when acetate and glucose are consumed simultaneously. On the other hand, acetate consumption after glucose depletion likely facilitates recovery following overshoot. Using acetate as an alternative carbon source when glucose is exhausted would further support the role of metabolic overflow in preventing ecological collapse and enhancing population resilience in fluctuating environments.

Acetate can also be toxic to bacterial cells, especially at high concentrations [6, 20]. Since our model focuses on second-order effects, it does not account for the increase in mortality caused by acetate toxicity, which directly affects the population size (see equation 4). In bacterial populations growing on glucose, high acetate levels (5-6 g/L) can reduce growth rates by up to 50% [32]. However, this effect is already noticeable at concentrations as low as 0.5 g/L [24], with significant reductions (approximately 20%) observed at 1 g/L [32]. While toxicity becomes significant at high concentrations, the substantial reductions in growth at lower levels suggest that regulation plays a key role in limiting growth before toxicity occurs. In any case, an increase in mortality due to high extracellular acetate concentrations would further help reduce overshoot, in line with the hypothesized role of overflow as a mechanism to prevent ecological collapse.

The idea of an active mechanism to reduce growth rates in microbial populations may seem counterintuitive. However, our results suggest that this effect of metabolic overflow can be crucial for the survival of microbial communities. From this perspective, overflow metabolism can be viewed as a cooperative strategy that reduces the risk of extinction. For this strategy to be effective, different species must cooperate by excreting and responding to a shared inhibitory signal.

Social behavior and collective decision-making are well-documented in both bacteria and yeasts [56, 57]. For instance, quorum sensing provides a mechanism for cell-to-cell communication, allowing microbial populations to adjust their behavior based on cell density [58, 59]. This process depends on the production of extracellular signals that convey information about cell density and trigger coordinated, population-wide responses.

Our results suggest that acetate functions as a quorum-sensing signal, providing information about global growth rates within bacterial communities (see Fig. 1). Furthermore, it helps prevent excessive growth by coordinating metabolic activity among coexisting populations. Ethanol and lactate would play similar roles in yeasts and mammal cells respectively. During exponential growth in nutrient-rich media, yeasts produce ethanol through fermentation, even in the presence of oxygen—a phenomenon known as the Crabtree effect [7]. Similar to acetate, ethanol is excreted into the surrounding environment, and its accumulation inhibits yeast growth. At low concentrations, this inhibition occurs through the increased expression of negative regulators of the G1/S transition, which prevents entry into the S phase and delays cell-cycle progression [14]. Ethanol is also toxic at high concentrations, reducing cell viability and increasing cell death [15, 18].

In mammalian cells, lactate production through fermentation under aerobic conditions, coupled with an increase in metabolic activity, is known as the Warburg effect [60]. At high concentrations, lactate causes acidification, which can be toxic to cells. Remarkably, artificially preventing lactic acidosis in tumors leads to rapid glucose exhaustion and population collapse [41]. In contrast, lactic acidosis allows tumor cells to maintain progressive proliferation for longer periods of time [41]. This suggests that metabolic overflow helps regulate cell growth and prevent collapse, in line with the predictions of our model. As occurs with acetate and ethanol, lactate can be viewed as conveying information about cellular metabolic within a tissue [10], explaining its role as a key regulator in multiple signalling pathways [61, 62].

The existence of a negative feedback loop between metabolic overflow and growth rates is well established. Our results suggest that this is not an accidental coincidence, but rather an active mechanism evolved to decelerate growth in nutrient-rich environments. This mechanism has profound implications for our understanding of microbial population dynamics, as it challenges the traditional view of microbial growth and highlights the existence of adaptive strategies to avoid overgrowth and collapse. In this regard, it has been suggested that the stationary phase in bacterial growth is not merely the passive result of resource depletion or toxin accumulation [63]. Instead, the transition into the stationary phase is an active and highly regulated process that involves both genetic and phenotypic changes [63].

This implies that bacterial cells not only transiently reduce their growth rates, but can also actively decide to stop growing even under favorable conditions. Acetate overflow could account for the regulated transition between exponential growth and stationary phase in bacterial populations. The entry into the stationary phase coincides with a marked increase in protein acetylation [64, 65], a key post-translational modification that regulates numerous central processes in bacterial cells, such as glycolysis [25], the TCA cycle, amino acid biosynthesis, or ribosome assembly [65, 66]. Acetate overflow and protein acetylation are intimately connected [19].

During exponential growth, acetate is produced from excess acetyl-CoA via acetyl-phosphate (AcP) [19, 67, 68], which is the primary source of acetyl groups for acetylation [26, 69]. Increased acetylation resulting from elevated AcP levels contributes to reducing growth rates [13] and developing stationary-phase phenotypes [23]. Acetylation is widespread across bacterial species [21–23] and it can be strongly induced by various carbon sources [68].

This suggests that acetate overflow might serve as a signal to trigger the transition from exponential growth to the stationary phase. Bacterial communities would not passively respond to resource depletion. Instead, they would actively regulate their proliferation rates, anticipating resource exhaustion and initiating the stationary phase to prevent ecological collapse. From this perspective, the carrying capacity would not be dictated solely by environmental constraints but would emerge from the interplay between these constraints and internal regulatory mechanisms designed to decelerate growth.

Our findings suggest that bacterial populations, through mechanisms like acetate overflow, are capable of self-regulating their growth to avoid overshoot and ecological collapse. This ability to prioritize long-term stability over short-term gains ensures the survival of microbial communities, even under competitive pressures. The fact that this strategy seems counterintuitive underscores how far we are from such a model in human population dynamics. Yet, the capacity to balance short-term needs with long-term sustainability is also a significant challenge for human societies [70].

## Code availability

Numerical simulations have been performed using Wolfram Mathematica. The code used in these simulations is available at the Notebook Archive (https://notebookarchive.org/2025-05-0h29lnh).

