## Supplementary Material A for "Metabolic overflow prevents ecological collapse in microbial populations"

### **Supplementary Information for:** Metabolic overflow prevents ecological collapse in microbial populations

[1]Grupo Interdisciplinar de Sistemas Complejos (GISC), 28040 Madrid, Spain.
[2]Departamento de Ecología, Universidad Complutense de Madrid, 28040 Madrid, Spain.
[3]Plasticentropy, 51100 Reims, France.
[4]Departamento de Inmunología, Universidad Complutense de Madrid, 28040 Madrid, Spain.

#### Supplementary Material A.
#### Effect of metabolic overflow in ecological competition.

This section provides the specification of the models used to generate the results shown in Figure 7 of the main text. To model the dynamics of two competing species, $P_1$ and $P_2$, equations 10 can be written as:

$$\begin{cases} P_1'(t) = g_1(t), \\ g_1'(t) = \rho_1\, e^{-\gamma_1\, A_1(t)} P_1(t) \left(1 - \dfrac{P_1(t)}{k_1 + \lambda_1\, R(t)}\right) - \mu_1\, P_1'(t), \\ P_2'(t) = g_2(t), \\ g_2'(t) = \rho_2\, e^{-\gamma_2\, A_2(t)} P_2(t) \left(1 - \dfrac{P_2(t)}{k_2 + \lambda_2\, R(t)}\right) - \mu_2\, P_2'(t), \\ R'(t) = \varphi - \delta_1\, P_1(t)R(t) - \delta_2\, P_2(t)R(t), \\ A'(t) = \alpha_1 \max\big(0, g_1(t) - \tilde{g}_1\big) + \alpha_2 \max\big(0, g_2(t) - \tilde{g}_2\big) \\ \qquad -\beta_1\, A(t)P_1(t) - \beta_2\, A(t)P_2(t), \end{cases} \quad \text{for } P_1 \geq 0 \text{ and } P_2 \geq 0.$$

with initial conditions

$$P_1(0) = P_{01}, g_1(0) = g_{01}, P_2(0) = P_{02}, g_2(0) = g_{02}, R(0) = R_0, A(0) = 0.$$

This system is integrated as long as both populations are viable, i.e., $P_1(t) > 0$ and $P_2(t) > 0$. If one of the populations goes extinct, the system transitions to a reduced version that excludes the extinct population. For example, if $P_2(t)$ reaches zero at time $t = t_e$, the system becomes:

$$\begin{cases} P_1'(t) = g_1(t), \\ g_1'(t) = \rho_1\, e^{-\gamma_1\, A_1(t)} P_1(t) \left(1 - \dfrac{P_1(t)}{k_1 + \lambda_1\, R(t)}\right) - \mu_1\, P_1'(t), \\ R'(t) = \varphi - \delta_1\, P_1(t)R(t), \\ A'(t) = \alpha_1 \max\big(0, g_1(t) - \tilde{g}_1\big) - \beta_1\, A(t)P_1(t), \end{cases} \quad \text{for } P_1 \geq 0.$$

with updated initial conditions at time $t = t_e$. The system is reduced analogously if $P_1(t)$ goes extinct instead.

Parameter values for each simulation are sampled uniformly within the ranges listed in Table S1. The outcome of each simulation is classified according to the long-term fate of the populations: coexistence (both populations survive), partial extinction (one of the populations disappears), or total extinction (both populations disappear).

| **Parameter** | **Min** | **Max** |
|---|---|---|
| $P_0$ | 0.1 | 1 |
| $R_0$ | 500 | 1000 |
| $\varphi$ | 0 | 10 |
| $\delta$ | 0.1 | 0.5 |
| $\beta$ | 0.1 | 0.3 |
| $k_i$ | 0.1 | 1 |
| $R_i$ | 0.1 | 0.5 |
| $\mu_i$ | 0.3 | 0.6 |
| $\lambda_i$ | 1 | 4 |
| $g_{0i}$ | 0 | 0.3 |
| $\alpha_i$ | 100 | 200 |
| $G_i$ | 0.1 | 0.5 |

**Table S1.** Parameter values used in Figure 7.

Figure 7 in the main text illustrates the outcomes of survival and extinction under three alternative scenarios regarding the use of acetate overflow:

- S1: Neither population uses acetate overflow
- S2: Only Population 1 uses acetate overflow
- S3: Both populations use acetate overflow

In the sections that follow, we provide a statistical comparison of the performance of acetate overflow under these scenarios. All analyses were performed using Mathematica, which offers robust tools for hypothesis testing. Specifically, the LocationEquivalenceTest function was used to determine the most appropriate statistical test for each dataset. This function adaptively selects the optimal method based on the underlying distributional properties, ensuring valid comparisons across scenarios.

Figure 7 comprises eight subpanels (A–H), covering both low and high mortality conditions, colonization dynamics, and responses to pulses of resources. For each panel, we summarize the survival probabilities or extinction outcomes under scenarios S1–S3, and provide statistical comparisons between them.

**Data shown in Figure 7 A. Colonization of a new environment (Low mortality)**

| Scenario | Population 1 survives | Population 2 survives |
|---|---|---|
| S1 | $41.089 \pm 0.613$ | $41.292 \pm 0.520$ |
| S2 | $83.476 \pm 0.300$ | $33.514 \pm 0.422$ |
| S3 | $80.096 \pm 0.301$ | $80.024 \pm 0.370$ |

**Table S2.** Mean ± Standard Deviation in Figure 7A.

| | Test | Statistic | P-Value |
|---|---|---|---|
| **S1 vs S2** | Kruskal-Wallis | 14.318 | $7.04602 \times 10^{-7}$ |
| **S1 vs S3** | Kruskal-Wallis | 14.2965 | $7.34814 \times 10^{-7}$ |
| **S2 vs S3** | K-Sample T | 630.996 | $1.82405 \times 10^{-15}$ |

**Table S3.** Pairwise comparison of survival of population 1.

| | Test | Statistic | P-Value |
|---|---|---|---|
| **S1 vs S2** | K-Sample T | 1349.72 | $2.20979\times10^{-18}$ |
| **S1 vs S3** | K-Sample T | 36881.3 | $2.90174\times10^{-31}$ |
| **S2 vs S3** | K-Sample T | 68665.6 | $1.08169\times10^{-33}$ |

**Table S4.** Statistical comparisons between survival scenarios S1, S2, and S3.

**Data shown in Figure 7 B. Survival and extinction of both populations during the colonization of a new environment (Low mortality)**

| Scenario | Both survive | Both collapse |
|---|---|---|
| S1 | $12.714 \pm 0.311$ | $30.333 \pm 0.410$ |
| S2 | $28.451 \pm 0.418$ | $11.461 \pm 0.302$ |
| S3 | $67.904 \pm 0.242$ | $7.784 \pm 0.300$ |

**Table S5.** Summary statistics (mean ± standard deviation) for the results shown in Figure 7B of the main text.

| | Test | Statistic | P-Value |
|---|---|---|---|
| S1 vs S2 | K-Sample T | 9116.33 | $8.31809\times10^{-26}$ |
| S1 vs S3 | K-Sample T | 196416. | $8.44871\times10^{-38}$ |
| S2 vs S3 | K-Sample T | 66730.1 | $1.39904\times10^{-33}$ |

**Table S6.** Statistical comparisons between survival scenarios S1, S2, and S3.

| | Test | Statistic | P-Value |
|---|---|---|---|
| S1 vs S2 | K-Sample T | 13742.2 | $2.08141\times10^{-27}$ |
| S1 vs S3 | K-Sample T | 19731. | $8.05371\times10^{-29}$ |
| S2 vs S3 | K-Sample T | 745.364 | $4.22474\times10^{-16}$ |

**Table S7.** Statistical comparisons between survival scenarios S1, S2, and S3.

**Data shown in Figure 7 C. Probability of survival of population 1 and 2 after after a pulse of resources in a community at steady state (Low mortality)**

| Scenario | Population 1 survives | Population 2 survives |
|---|---|---|
| S1 | $24.709 \pm 0.448193$ | $24.662 \pm 0.357174$ |
| S2 | $67.175 \pm 0.530079$ | $14.065 \pm 0.35246$ |
| S3 | $42.096 \pm 0.292582$ | $42.136 \pm 0.415751$ |

**Table S8.** Summary statistics (mean ± standard deviation) for the results shown in the analysis of three strategies and two competing populations.

| | Test | Statistic | P-Value |
|---|---|---|---|
| S1 vs S2 | K-Sample T | 37425. | $2.54382\times10^{-31}$ |
| S1 vs S3 | K-Sample T | 10552.5 | $2.2347\times10^{-26}$ |
| S2 vs S3 | K-Sample T | 17157. | $2.83043\times10^{-28}$ |

**Table S9.** Statistical comparisons between survival scenarios S1, S2, and S3.

| | Test | Statistic | P-Value |
|---|---|---|---|
| S1 vs S2 | K-Sample T | 4459.73 | $5.09327\times10^{-23}$ |
| S1 vs S3 | K-Sample T | 10163.7 | $3.13101\times10^{-26}$ |
| S2 vs S3 | K-Sample T | 26524.5 | $5.62838\times10^{-30}$ |

**Table S10.** Statistical comparison results for different scenarios using the K-Sample T test.

**Data shown in Figure 7 D. Survival and extinction of both populations at steady state (Low mortality)**

| Scenario | Both populations survive | Both populations collapse |
|---|---|---|
| S1 | $10.298 \pm 0.223746$ | $60.927 \pm 0.457628$ |
| S2 | $12.235 \pm 0.314475$ | $30.995 \pm 0.592626$ |
| S3 | $22.642 \pm 0.396171$ | $38.41 \pm 0.334332$ |

**Table S11.** Summary statistics (mean ± standard deviation) for the results shown in the analysis of three strategies and two competing populations.

| | Test | Statistic | P-Value |
|---|---|---|---|
| S1 vs S2 | K-Sample T | 251.883 | $4.99566\times10^{-12}$ |
| S1 vs S3 | K-Sample T | 7360.6 | $5.68115\times10^{-25}$ |
| S2 vs S3 | K-Sample T | 4233.24 | $8.12668\times10^{-23}$ |

**Table S12.** Statistical comparison results for different scenarios using the K-Sample T test.

| | Test | Statistic | P-Value |
|---|---|---|---|
| S1 vs S2 | K-Sample T | 15980.7 | $5.36023\times10^{-28}$ |
| S1 vs S3 | K-Sample T | 15785.0 | $5.98824\times10^{-28}$ |
| S2 vs S3 | K-Sample T | 1187.56 | $6.88579\times10^{-18}$ |

**Table S13.** Statistical comparison results for different scenarios using the K-Sample T test.

**Data shown in Figure 7 E. Colonization of a new environment (High mortality)**

| Scenario | Population 1 survives | Population 2 survives |
|---|---|---|
| S1 | $76.628 \pm 0.323824$ | $76.613 \pm 0.348905$ |
| S2 | $94.595 \pm 0.222024$ | $75.732 \pm 0.405183$ |
| S3 | $94.13 \pm 0.203415$ | $94.077 \pm 0.16879$ |

**Table S14.** Summary statistics (mean ± standard deviation) for the results shown in the analysis of three strategies and two competing populations.

| | Test | Statistic | P-Value |
|---|---|---|---|
| S1 vs S2 | K-Sample T | 20940.6 | $4.71669\times10^{-29}$ |
| S1 vs S3 | K-Sample T | 20946.4 | $4.70495\times10^{-29}$ |
| S2 vs S3 | K-Sample T | 23.8469 | 0.000119588 |

**Table S15.** Statistical comparison results for different scenarios using the K-Sample T test.

| | Test | Statistic | P-Value |
|---|---|---|---|
| S1 vs S2 | K-Sample T | 27.1473 | 0.0000590259 |
| S1 vs S3 | Kruskal-Wallis | 14.3072 | $7.19578\times10^{-7}$ |
| S2 vs S3 | Kruskal-Wallis | 14.3072 | $7.19578\times10^{-7}$ |

**Table S16.** Statistical comparison results for different scenarios using the K-Sample T test and Kruskal-Wallis test.

**Data shown in Figure 7 F. Survival and extinction of both populations during the colonization of a new environment (High mortality)**

| Scenario | Both populations survive | Both populations collapse |
|---|---|---|
| S1 | $61.295 \pm 0.336889$ | $8.054 \pm 0.151745$ |
| S2 | $73.864 \pm 0.398809$ | $3.537 \pm 0.178204$ |
| S3 | $90.105 \pm 0.27456$ | $1.898 \pm 0.108095$ |

**Table S17.** Summary statistics (mean ± standard deviation) for the results shown in the analysis of three strategies and two competing populations.

| | Test | Statistic | P-Value |
|---|---|---|---|
| S1 vs S2 | K-Sample T | 5796.5 | $4.84978 \times 10^{-24}$ |
| S1 vs S3 | K-Sample T | 43944.6 | $5.99869 \times 10^{-32}$ |
| S2 vs S3 | K-Sample T | 11251.4 | $1.25579 \times 10^{-26}$ |

**Table S18.** Statistical comparison results for different scenarios using the K-Sample T test.

| | Test | Statistic | P-Value |
|---|---|---|---|
| S1 vs S2 | K-Sample T | 3724.36 | $2.5607 \times 10^{-22}$ |
| S1 vs S3 | K-Sample T | 10917.6 | $1.64608 \times 10^{-26}$ |
| S2 vs S3 | K-Sample T | 618.382 | $2.17704 \times 10^{-15}$ |

**Table S19.** Statistical comparison results for different scenarios using the K-Sample T test.

**Data shown in Figure 7 G. Probability of survival of population 1 and 2 after after a pulse of resources in a community at steady state (High mortality)**

| Scenario | Population 1 survives | Population 2 survives |
|---|---|---|
| S1 | $62.095 \pm 0.332374$ | $61.834 \pm 0.515562$ |
| S2 | $90.073 \pm 0.225095$ | $57.463 \pm 0.488127$ |
| S3 | $85.493 \pm 0.262173$ | $85.561 \pm 0.58108$ |

**Table S20.** Summary statistics (mean ± standard deviation) for the results shown in the analysis of three strategies and two competing populations.

| | Test | Statistic | P-Value |
|---|---|---|---|
| S1 vs S2 | K-Sample T | 48576.9 | $2.43491\times10^{-32}$ |
| S1 vs S3 | K-Sample T | 30549.4 | $1.57947\times10^{-30}$ |
| S2 vs S3 | K-Sample T | 1756.78 | $2.11547\times10^{-19}$ |

**Table S21.** Statistical comparison results for different scenarios using the K-Sample T test.

| | Test | Statistic | P-Value |
|---|---|---|---|
| S1 vs S2 | K-Sample T | 379.026 | $1.53252\times10^{-13}$ |
| S1 vs S3 | K-Sample T | 9329.06 | $6.76127\times10^{-26}$ |
| S2 vs S3 | K-Sample T | 13708.4 | $2.128\times10^{-27}$ |

**Table S22.** Statistical comparison results for different scenarios using the K-Sample T test.

**Data shown in Figure 7 H. Survival and extinction of both populations at steady state (High mortality)**

| Scenario | Both populations survive | Both populations collapse |
|---|---|---|
| S1 | $48.568 \pm 0.422869$ | $24.639 \pm 0.433985$ |
| S2 | $56.46 \pm 0.486438$ | $8.924 \pm 0.227947$ |
| S3 | $78.172 \pm 0.464753$ | $7.118 \pm 0.320062$ |

**Table S23.** Summary statistics (mean ± standard deviation) for the results shown in the analysis of three strategies and two competing populations.

| | Test | Statistic | P-Value |
|---|---|---|---|
| S1 vs S2 | K-Sample T | 1499.22 | $8.68196\times10^{-19}$ |
| S1 vs S3 | K-Sample T | 22197.8 | $2.79214\times10^{-29}$ |
| S2 vs S3 | K-Sample T | 10415.2 | $2.51364\times10^{-26}$ |

**Table S24.** Statistical comparison results for different scenarios using the K-Sample T test.

| | Test | Statistic | P-Value |
|---|---|---|---|
| S1 vs S2 | K-Sample T | 10277.1 | $2.83406\times10^{-26}$ |
| S1 vs S3 | K-Sample T | 10557.2 | $2.2257\times10^{-26}$ |
| S2 vs S3 | K-Sample T | 211.246 | $2.18242\times10^{-11}$ |

**Table S25.** Statistical comparison results for different scenarios using the K-Sample T test.
